# Proteomic analysis of gut in *Labeo rohita* reveals ECM as Key Player in host’s Response to *Aeromonas hydrophila* Infection

**DOI:** 10.1101/2024.02.19.581092

**Authors:** Mehar Un Nissa, Nevil Pinto, Biplab Ghosh, Anwesha Banerjee, Urvi Singh, Mukunda Goswami, Sanjeeva Srivastava

## Abstract

In the aquaculture sector, one of the challenges include disease outbreaks such as bacterial infections, particularly from *Aeromonas hydrophila* (*Ah*), impacting both wild and farmed fish. In this study, we conducted a proteomic analysis of the gut tissue in *Labeo rohita* following *Ah* infection to elucidate the protein alterations and its implications for immune response. Our findings reveal significant dysregulation in extracellular matrix (ECM) associated proteins during *Ah* infection, with increased abundance of elastin and Collagen alpha-3(VI) contributing to matrix rigidity. Pathway and enrichment analysis of differentially expressed proteins (DEPs) highlights the involvement of ECM-related pathways, including Focal adhesions, Integrin cell surface interactions, and actin cytoskeleton organization.

Focal adhesions, crucial for connecting intracellular actin bundles to the ECM, play a pivotal role in immune response during infections. Increased abundance of integrin alpha 1, integrin beta 1, and Tetraspanin suggests their involvement in the host’s response to *Ah* infection. Proteins associated with actin cytoskeleton reorganization, such as myosin, tropomyosin, and phosphoglucomutase, exhibit increased abundance, influencing changes in cell behavior. Additionally, upregulated proteins like LTBP1 and Fibrillin-2 contribute to TGF-β signaling and focal adhesion, indicating their role in immune regulation.

The study also identifies elevated levels of laminin, galectin 3, and tenascin-C, which interact with integrins and other ECM components, influencing immune cell migration and function. These proteins, along with decorin and lumican, act as immunomodulators, coordinating pro- and anti-inflammatory responses. ECM fragments released during pathogen invasion serve as “danger signals,” initiating pathogen clearance and tissue repair through Toll-like receptor signaling.

**IMPORTANCE:** The study underscores the critical role of the extracellular matrix (ECM) and its associated proteins in the immune response of aquatic organisms during bacterial infections like *Aeromonas hydrophila* (Ah). Understanding the intricate interplay between ECM alterations and immune response pathways provides crucial insights for developing effective disease control strategies in aquaculture. By identifying key proteins and pathways involved in host defense mechanisms, this research lays the groundwork for targeted interventions to mitigate the impact of bacterial infections on fish health and aquaculture production.

## INTRODUCTION

Aquaculture stands out as one of the most rapidly expanding sectors, serving as a crucial source of high-quality protein to ensure global food security and foster international trade. However, the intensive aquaculture sector faces challenges such as disease emergence and stress factors, demanding careful consideration for sustainable production goals. Fish frequently encounter pathogens in environment comprising viruses, bacteria, fungi, and parasites. The primary disease causing bacteria in carps belong to Aeromonas spp., a highly opportunistic pathogen accounting for 66.66% of bacterial infections in many fish types including carps, salmon, trout (1).

Bacterial infections cause high mortality rates in both wild and farmed fish (2). The strains encountered maximally are *A. hydrophila*, *A. caviae*, *A. veronii*, A. *salmonicida* and *A. sobria*. Among these bacteria, *A. hydrophila* has been considered as the most harmful pathogen to aquatic species, mainly causing haemorrhagic disease in fish farms (3). Since it’s difficult to screen them directly from natural resources, these do not get detected until diagnosis of symptoms associated with diseases like aeromonad septicemia/ bacteremia and gastroenteritis are observed. The oral route serves as the predominant mode of entry, exposing the gut to potential infections. Successful manifestation of disease requires bacteria to adhere, invade, and colonize the host, evading immune responses through various mechanisms and substrates (4, 5). The tissue specificity of bacterial invasion, particularly in the gut, underscores the critical role of the extracellular matrix (ECM) as a facilitator of infection. The ECM, a fundamental cellular component, not only provides structural support but also plays a vital signaling role (6). It creates a microenvironment for essential signals that govern immune cell migration, activation, proliferation, and differentiation within infected tissues (7). Each marked tissue-specific pathogenic infection has a distinct ECM signature. Different tissue-specific infections exhibit distinct ECM signatures, comprising structural matrix components like collagen, proteoglycans, glycosaminoglycans, and functional elements such as enzymes and growth factors (8). The enzymes repair and remodel the structural components by enhanced synthesis and deposition, degradation and protein modification, ultimately impacting the overall micro-environment essential for the infection and inflammatory response. The breakdown of primary and secondary defenses by bacterial enzymes, such as collagenases and hyaluronidases, significantly affects overall immunity (9).

A peek into the dynamics of the unidentified multifactorial pathogenesis of this bacterium is offered by omics studies ranging from genomics, transcriptomics and proteomics, enabling better understanding and development of effective disease control strategies. Lie Jin et. al have constructed the complete genome sequence of virulent strain, *A. hydrophila* HX-3 and drawn its comparison to other strains in order to identify key genes responsible for virulence and quorum sensing (10). Another team, employing BOX-PCR-based DNA fingerprinting evaluated banding patterns to investigate the strain level genotypic markers in *Ah* (11). The study by Hu et al, delve through transcriptomics and proteomics to identify mucosal immunity i.e. first line of defence in gills of *Carassius auratus* against *Ah* (12). With the tangent of liver tissue analysis, a proteomic study suggests that *Ah* infection modulates host metabolic pathways and innate immunity with identified markers such as Toll like receptors and C-lectins in *Labeo rohita* (13). Our study builds upon these efforts by employing holistic discovery-based and validation-based proteomic methods to unravel the pathogenesis of *A. hydrophila* in the gut tissue of *Labeo rohita (rohu)*. This species is a significant Indian carp, known for its nutritional value and widespread consumer preference, and ranks among the top ten inland fish produced globally (14). We have provided an overall analysis of the relative changes observed in protein profiles pertaining to gut tissue and have narrowed our findings to ECM-dominant inferences. This has enabled us to identify pathways and markers that if perturbed can be used for both disease diagnosis and treatment of clinical manifestations.

## MATERIALS AND METHODS

### Overall experimental design

This study involved a proteomic analysis of gut tissue in *Labeo rohita* following infection with *Aeromonas hydrophila (Ah)*. Fish with an average weight of 70±10 g and length of 19±1 cm were infected with *Ah* and then sampled, resulting in a total of 12 samples, with six from the Control group and six from the *Ah* infected group (AH). From these samples, four from each group were selected for discovery-based proteomic analysis. The MaxQuant software was used to analyze the data, followed by statistical analysis in the Metaboanalyst tool to identify the differentially expressed proteins (DEPs; p value 0.05, fold change 1.5). An overview of the functional annotation of altered proteins and metabolic pathways was obtained by gene ontology (GO) analysis. The SRM technique was used to validate the protein abundance changes for a panel of differentially expressed proteins using all 12 samples.

### Maintenance of fish for the experiment

Six-month-old fish (N=150; average weight 70±10 g) were obtained from a nearby fish farm in the Pen Raigad District of Maharashtra and transported to a wet lab facility (ICAR-CIFE, Mumbai) for the study. In three circular fibre tanks, the fish were evenly spaced, acclimated, and fed 2% of their body weight. The fish were kept in a temperature range of 26 to 28 °C with adequate aeration and daily faeces removal. Every fish was checked for external clinical symptoms, and a random subset of fish from each of the tanks was slaughtered in order to look for pathogens in the fish using PCR.

### Bacterial challenge and tissue sampling

Bacterial isolation and identification was done as reported in our previous work (13). In brief, tissue from a naturally co-infected fish was used for isolation of *Ah* strain (NCBI Accession no. MT374248) which was confirmed using PCR. Fishes were distributed into six high density polyethylene plastic crates, each holding six fish and kept at 26-28 °C. Three crates labelled as AH group were intraperitoneally inoculated with LD_50_ dose of 1.5×10^8^ bacterial cells in PBS. Three crates of Control group were subjected to equivalent volume of PBS injection. Following infection, there were visible indications of haemorrhage in the AH group. As would be expected, the control group did not exhibit these symptoms. Gut samples were collected 48 hours post challenge and stored at -80 °C till further use. Tissue from three fish was combined into one yielding a total of six samples each for the Control and AH groups. The samples were designated as Gut-C1 to C6 and Gut-AH1 to AH6, respectively.

### Tissue lysate preparation and protein extracts

SDS-containing lysis buffer (5% SDS, 100mM Tris/HCl pH 8.5 (adjusted with phosphoric acid)) was used to prepare the tissue lysates. The tissue was weighed (40–50 mg) and washed in a 1X phosphate buffered saline (PBS) solution twice–thrice. Following it, 250 µl of lysis buffer and 5 µl of protease inhibitor cocktail (50X stock, Sigma-Catalogue no. 11873580001) were added to the tissue. It was then left to sit on ice for 30 minutes. Sonication was done for two minutes at a 40% amplitude, and pulse 5 sec on/off. The debris was removed using centrifugation, and the clear supernatant was collected.

### Quantification and digestion of proteins

Tissue protein quantification and digestion were done as mentioned in our previous work (15). In brief, protein was quantified through BCA method using Bovine serum Albumin as standard. Before digestion, 30 µg of protein (in SDS lysis buffer) was reduced with 20mM TCEP. Reduced protein was loaded onto a 30KDa column for downstream steps of alkylation and digestion. Iodoacetamide was used for alkylation. For digestion, trypsin in 50mM ABC in 1:30 ratio with protein was added over the column itself and kept in wet chamber at 37 °C for 16 hours. Peptides were dried and stored. Before mass spectrometry analysis, peptides were cleaned using in-house prepared C18 tips (66883-U-Merck).

### Data acquisition with liquid chromatography tandem mass spectrometry

Peptides were quantified for each sample using the Scopes method (16). Following quantification, 1µg of the peptide sample was injected to the mass spectrometer for LC-MS/MS analysis. Eight samples were run with a 120-minute LC gradient with a flow rate of 300 nl/min. Mobile phase included solvent A as 0.1% Formic acid (FA) and solvent B as 80% Acetonitrile (ACN) with 0.1% FA. Samples included four each of the Control and AH groups, respectively. An Easy-nLC nano-flow liquid chromatography 1200 system linked with an Orbitrap-Fusion Tribrid mass spectrometer, was used to generate the data in data dependent acquisition mode. Peptides were loaded onto the trap column (Thermo Fisher Scientific, 100 mm x 2 cm, nanoViper C18, 5 mm, 100A) at a flow rate of 5 µl/min and then passed through the analytical column (Thermo Fisher Scientific, 75 μm × 50 cm, 3 μm particle, and 100 Å pore size). MS1 parameters included Orbitrap resolution 60k, scan range 375-1700, RF Lens 60%, maximum injection time 50s, exclusion duration 40s and mass tolerance 10 ppm. MS2 settings were isolation mode Quadrupole, isolation window 2, activation type HCD, Orbitrap resolution 15k, maximum injection time 30ms, data type centroid. For MS1 and MS2 level, the AGC target was set at 400000 and 10000, respectively. For positive internal calibration, a lock mass of 445.12003 m/z was used.

### Protein identification and label-free quantification

The raw data obtained from LC/MS-MS was searched in MaxQuant (v1.6.6.0) software through Andromeda search engine. Under group specific parameters, multiplicity was set as standard and label as 1. For modifications, oxidation at Methionine (+15.994915 Da) and Acetyl (protein N term) were chosen. Carbamidomethyl (C) was set under fixed modifications. Data was analysed in Label free quantification mode and match between runs was enabled. Trypsin/P was chosen as protease and maximum of two missed cleavages were allowed. Instrument type was selected as Orbitrap keeping all parameters as default. Under global parameters tab, *Labeo rohita* UniProt whole protein sequences (UP000290572, ID-84645) database was selected and include contaminant option was enabled. Minimum peptide length was set to 7. Maximum precursor mass tolerances were set at 20 ppm in the first search, 4.5 ppm in the main search, and 20 ppm for fragment mass tolerances. The false discovery rate for proteins, peptides, and PSMs was 1%. Proteins were only recognized by their unique peptide, and reverse was chosen as the decoy mode option. For a quantitative comparison, the LFQ intensities obtained for each sample were taken into consideration (Table S1).

### Statistical analysis using MetaboAnalyst

Statistical analysis was performed taking MaxQuant output files using Metaboanalyst software (17). After MaxQuant analysis, the dataset comprised the identified proteins, along with LFQ intensities for 4 samples each from Control and AH groups. The data was filtered to remove features matched with contaminant or reverse sequences. The missing value imputation was done separately for Control and AH group using KNN feature wise option, and features with more than 30% missing values were removed. Further statistical analysis was performed taking the common features between Control and AH groups. No data filtration was applied and data was log10 transformed before being plotted. The fold change threshold was taken to be 1.5, and RAW p value cut off was 0.05 to get the DEPs (Table S1). Computation of the Variable importance in projection (VIP) score was done via the Partial Least Squares Discriminant Analysis (PLS-DA) which is a supervised method. As a weighted sum of squares of the PLS weight, VIP indicates the importance of the variable to the whole model. Heatmap representing the expression of top DEPs was also obtained from Metaboanalyst analysis. Volcano plots were plotted using online tool VolcanoseR(18). These significant DEPs were further considered for validation with reference to existing literature on their functions and roles in microbial diseases.

### Gene ontology, Pathway and protein-protein interaction enrichment analysis

Significant DEPs were considered for functional annotation and biological pathway analysis. Gene names of dysregulated proteins were retrieved from EggNOG (19) resource on the basis of ortholog annotation and from literature (as the gene names for *L. rohita* are not updated yet in the available databases). Metascape tool was used to obtain statistically enriched terms (GO/KEGG terms, canonical pathways) for all DEPs. STRING tool version 11.5 was used for protein-protein interaction (PPI) and visualization for the selected list of proteins from interested GO/KEGG terms/ canonical pathways (Table S2). Minimum overlap of 3, p-value cut off of 0.05, minimum enrichment 1.5 was employed.

### Validation of differential protein abundance using targeted proteomics

The proteomic validation was done using selected/ multiple reaction monitoring (SRM/MRM) method. The SRM method was created using transition list obtained from Skyline version 23.0.9.187. The Skyline setting included missed cleavage as 0, precursor charges +2, +3, and product charge was set at +1 with ‘y’ ion transitions (from ion 2 to last ion −1). Initial optimization was started with 12 upregulated proteins where a pooled peptide sample was run against six transition lists each with 350-400 transitions. For data acquisition, TSQ Altis mass spectrometer (ThermoFisher Scientific, USA) coupled to an HPLC-Dionex Ultimate 3000 system (ThermoFisher Scientific, USA) was utilised. Hypersil Gold C18 (1.9 μm, 100 × 2.1 mm, ThermoFisher Scientific, USA) reverse-phase column was used for peptide separation. A binary buffer system with flowrate at 0.45 ml/min was employed for 10 mins with buffer A and B constituting 0.1% FA and 80% ACN in 0.1% FA, respectively. For final runs, samples were spiked-in with an equal amount of heavy R labelled synthetic peptide ENQTCDIYNGEGR to ensure consistency across the runs. Including 10 transitions for synthetic peptide, the final method contained 402 transitions (Table S3). One μg of peptides from 12 samples, 6 each of Control and AH group was injected against the final method.

### Targeted proteomics data analysis

The result files (.raw) were imported to Skyline software and assigned to conditions of Control and AH, respectively. The DDA spectral library for analysis of SRM data was built using MSMS (.msms) file obtained from MaxQuant analysis. Data refinement was manually carried out considering the peak shape, retention time alignment, and dot product-based match with spectral library. The peptides not adhering to norms and specifications were deleted in order to refine the data. Statistical analysis was performed using MSstats external tool inbuilt in Skyline to identify peptides with significant fold change (cut off 1.5) and *P* value (≤ 0.05). Result reports for all the peptides containing their peak area values were exported in .csv format to carry out the further analysis (Table S3).

### Ethics statement

For this work, all fish were collected and sacrificed at The Indian Council of Agricultural Research, Central Institute of fisheries Education (ICAR-CIFE), Versova, Mumbai, 400076. The presented work is a part of sanctioned project of Department of Biotechnology, India (BT/PR15285/AAQ/3/753/2015) and the work was approved by Institute Ethical committee, ICAR-CIFE (Project code 1008979).

## RESULTS

### Differential gut proteome between controls and Ah infected group and overall functional annotation

Discovery based proteomics data was acquired for eight samples to derive a comparison of proteomes between Control and AH (48-hour post *Ah* infection). Label free quantification was used to compare the proteome profiles between infected and uninfected gut tissues (details in Material and Methods section and workflow shown in Fig. 1). The LFQ analysis resulted in the identification of 2632 proteins (features). After missing value imputation, the processed data included 1961 and 1744 as respective features for Control and AH with 1638 features accounting for the overlap (Table S1). Statistical analysis of the common features resulted in the identification of 148 DEPs of which 74 were upregulated and 74 were down regulated in AH group as compared to Control (Fig. 2A, Table S1). Using PLS-DA model based on VIP score, some features could distinguish AH from Control group. Fig. 2B shows top 20 features classifying the AH and Control group based on VIP scores. Further the samples were clearly clustered with similar abundance profiles using unsupervised hierarchical clustering which also stratified the *Ah* infected and Control groups as represented for top 25 features (Fig. 2C). A few of the DEPs including Tetraspanin (tspan), Elastin (eln), and Annexin (Anx) among upregulated are shown (Fig. 2D-F) and, Antithrombin-III (Serpinc1), and Hyaluronan-binding 2 (Habp2) and Peroxiredoxin-like 2 (Prxl2a) for downregulated proteins (Fig. 2G-I) in the AH group compared to Control.

**Figure 1.**
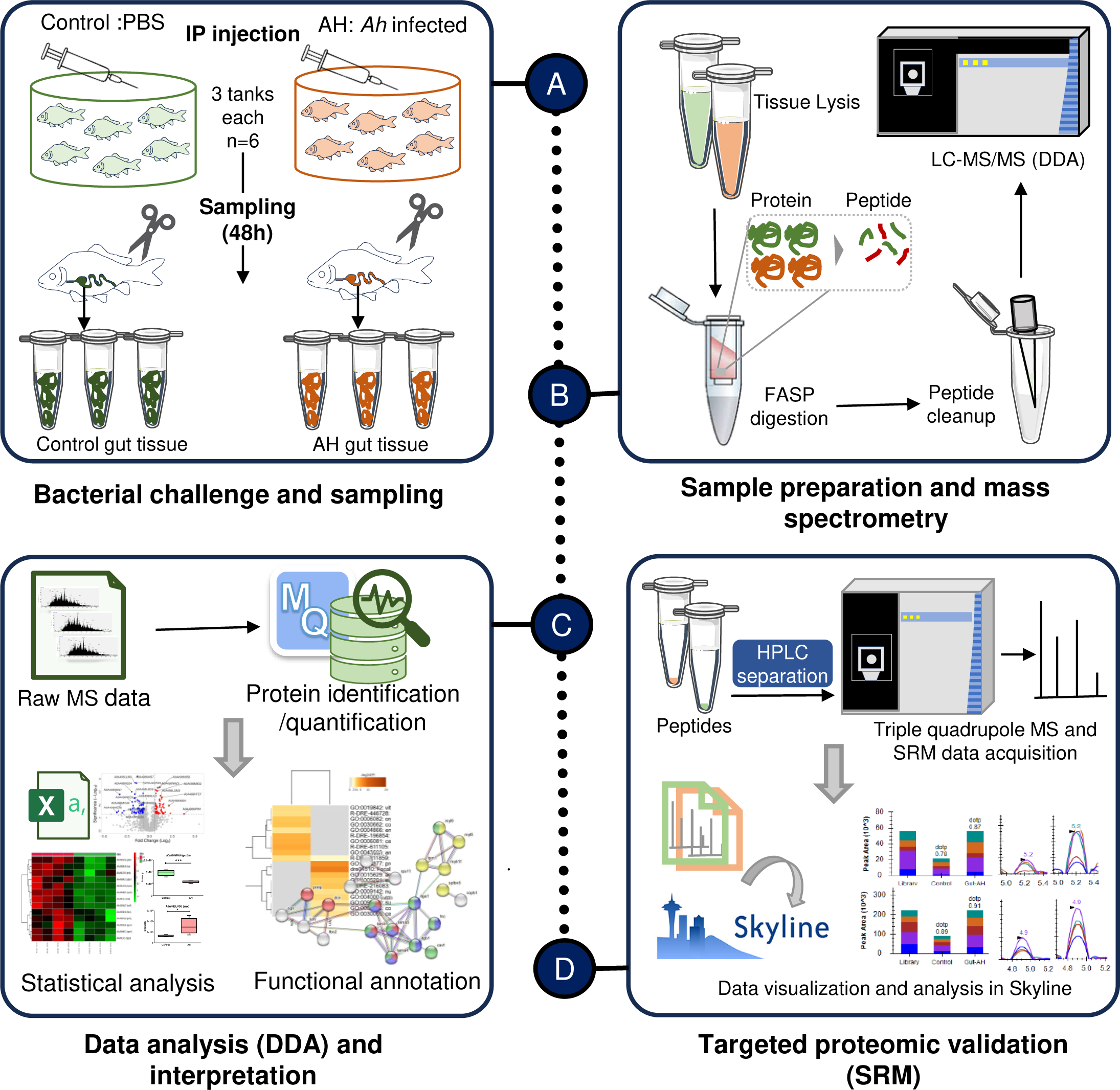
Schematic representation of the study: **(A)** In the challenge study, the Control group received intraperitoneal injections of Phosphate Buffer Saline (PBS), while the AH group (*Ah* challenged) received *A. hydrophila* injections. After 48 hours of infection, fish were euthanized, and gut samples were collected, as described in the text. **(B)** For proteomics analysis, tissues were lysed in a Sodium Dodecyl Sulphate (SDS)-containing buffer, and protein digestion was carried out using the filter-assisted sample preparation (FASP) method. Subsequently, peptide samples underwent cleaning before being subjected to mass spectrometry for Data-Dependent Acquisition (DDA) using Orbitrap mass spectrometry. **(C)** The acquired raw mass spectrometry data (.raw) underwent analysis with MaxQuant software for protein identification and quantification. Statistical analysis was performed to identify differentially expressed proteins, followed by functional analysis. **(D)** Targeted proteomic validation of selected proteins through selected reaction monitoring involved subjecting peptide samples to High-Performance Liquid Chromatography (HPLC). This was followed by target precursor and transition selection in the Triple Quadrupole mass spectrometer for the acquisition of spectral data. The Selected Reaction Monitoring (SRM) data were analyzed using Skyline software, and the targeted data were compared with the Spectral Library.

**Figure 2.**
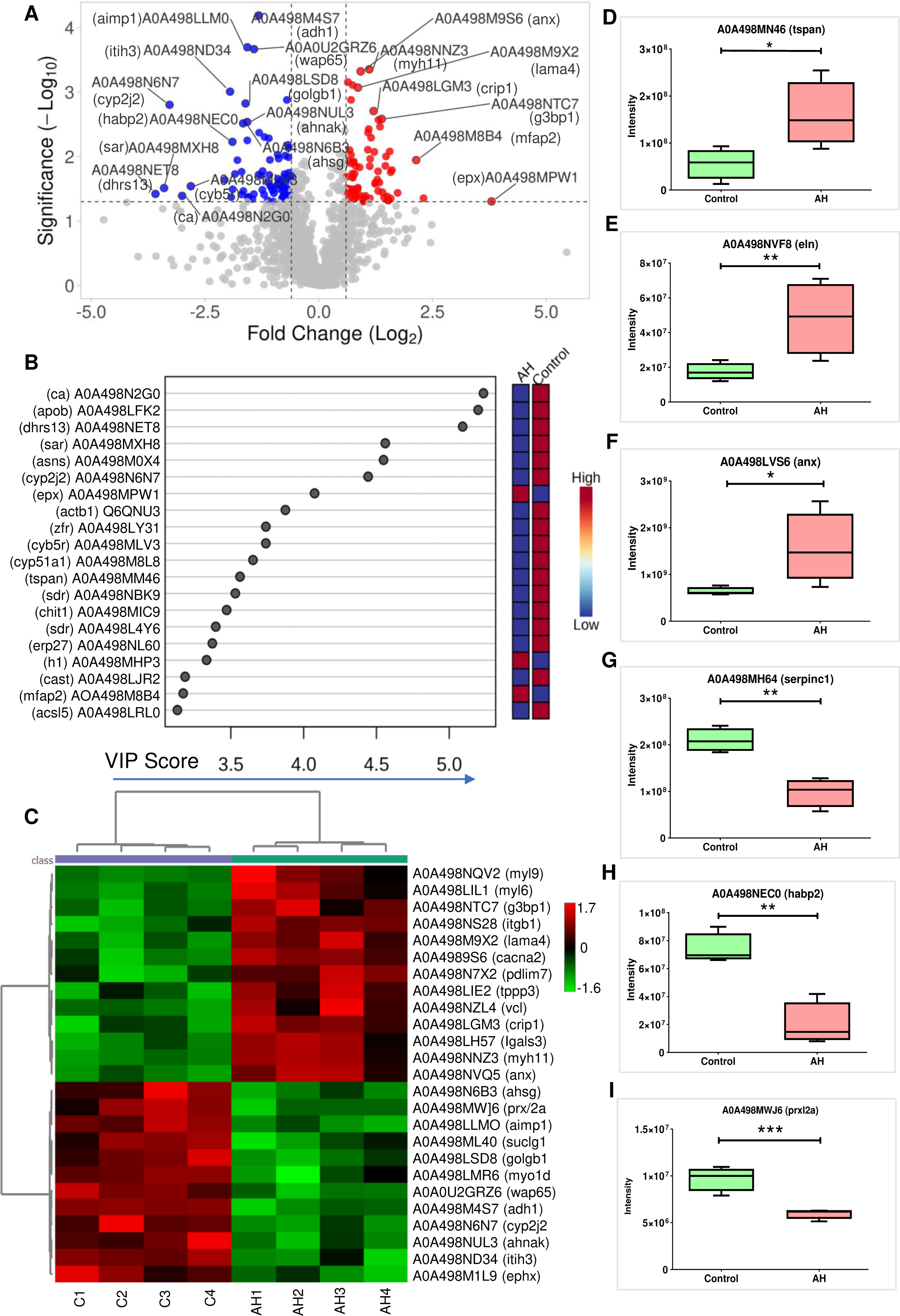
Shotgun Proteomic analysis reveals altered host gut proteome during *Ah* infection. **(A)**. Volcano plot depicting significantly altered protein candidates (Fold Change of 1.5, p <0.05), where red colour and blue representing upregulated and downregulated proteins respectively. (**B)** Top 20 altered key proteins found on the basis of VIP Score. **(C)** Heatmaps showing differential expression (abundances) of top 25 significant proteins (t-test, p ≤ 0.05) across 4 replicates each of control tissues (C1 to C4) and AH infected tissues (AH1 to AH4). D-I. Box plots representing altered abundances of 6 proteins (3 upregulated and 3 downregulated) in *Ah* infected group as compared to Control. (For Boxplots, *p ≤ 0.05, **p ≤ 0.01 and, ***p ≤ 0.001).

Functional mapping of upregulated and downregulated DEPs was done using Metascape to understand how AH infection influences on the host proteome qualitatively. Fig. 3A shows the significant GO terms and pathways mapped to the DEPs. For downregulated DEPs, the mapped GO terms include vitamin binding, organic acid metabolic process, endopeptidase inhibitor activity, cellular aldehyde and amide metabolic process. The mapped pathways include cell junction organisation, metabolism of vitamins and cofactors and respiratory electron transport. On the other hand, GO/pathways mapped to upregulated DEPs include protein containing complex binding, actin cytoskeleton, extracellular matrix structural constituent, supramolecular complex, focal adhesion, cell substrate junction and integrin cell surface interactions (Fig. 3A, Table S2).

**Figure 3.**
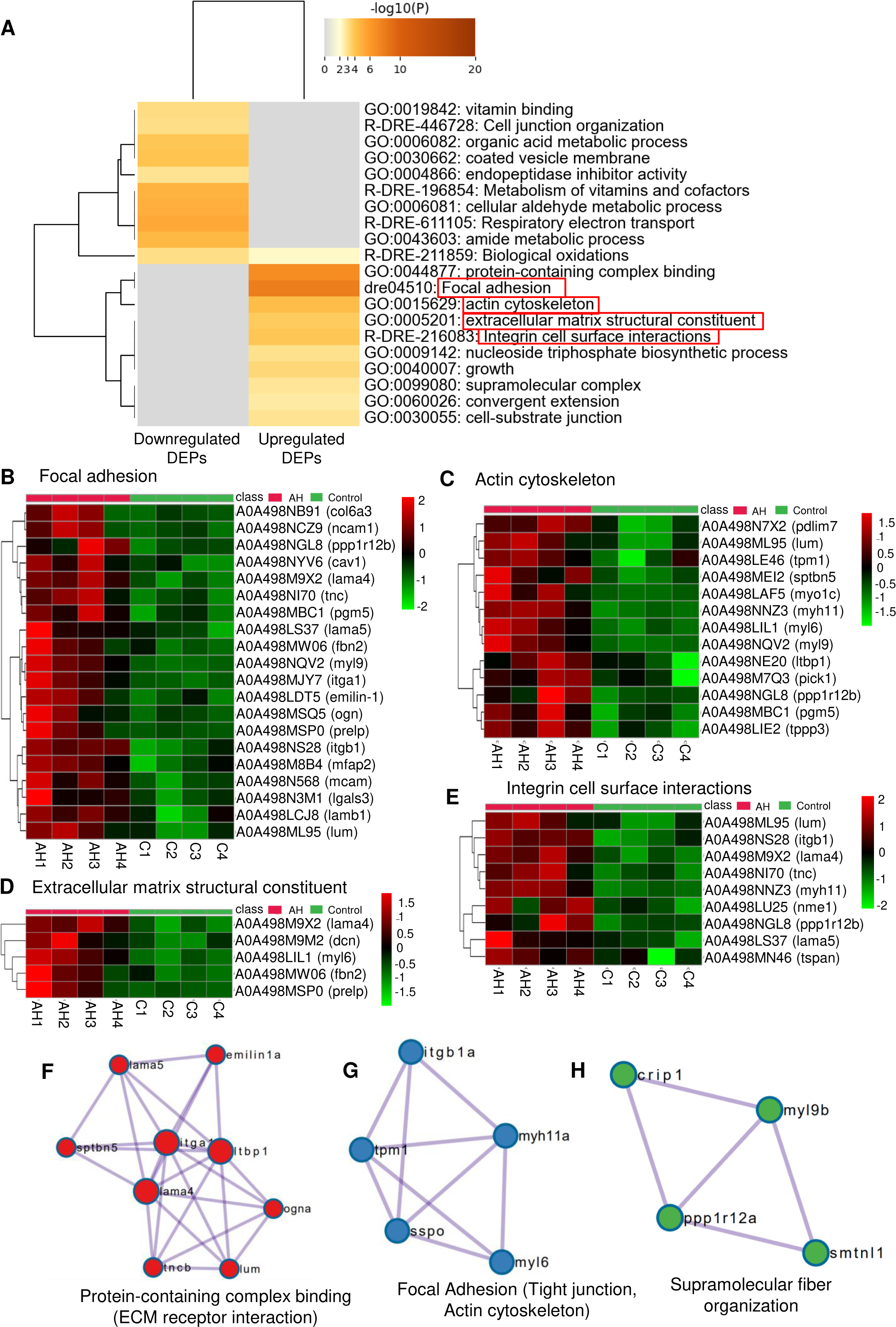
GO and pathway analysis for differentially expressed proteins. **(A**) Hierarchical cluster of significant GO/KEGG terms based on Kappa-statistical similarities among their gene memberships for upregulated and downregulated DEPs. Then 0.3 kappa score was applied as the threshold to cast the tree into term clusters. The term with the best p-value within each cluster as its representative term is displayed in a dendrogram. The heatmap cells are colored by their p-values, white cells indicate the lack of enrichment for that term in the corresponding gene list. Important pathways related to ECM are highlighted with red rectangle. **(B-E)** Heatmaps showing differential abundances of proteins mapped to Focal adhesion pathway, actin cytoskeleton, extracellular matrix structural constituent and integrin cell surface interactions pathway, respectively. **(F-H)** Top three Molecular Complex Detection (MCODE) network components for DEPs viz. protein-containing complex binding, Focal adhesion, supramolecular fiber organization.

### Ah infection altered proteins related to Focal adhesion and extracellular matrix pathways

As mentioned above, along with other pathways, enriched GO/pathways related to extracellular matrix (ECM) were Focal adhesion, actin cytoskeleton, Integrin cell surface interactions and extracellular matrix structural constituent. Enrichment of these pathways with high confidence is an indication that ECM in gut plays an important role during *Ah* infection. To focal adhesion pathway, 20 DEPs were mapped and similarly 13 proteins in actin cytoskeleton, 5 in the extracellular matrix constituent and, 9 in integrin cell surface interaction pathway were mapped (Fig. 3B-E). In total, 39 of unique significant upregulated DEPs were mapped to these processes/pathways highlighting the active focal adhesion and ECM signalling pathways in enterocytes. Further, in the Metascape tool, Molecular Complex Detection (MCODE) algorithm was applied on the enrichment network to identify neighborhoods where proteins are densely connected. Top MCODE network components for DEPs include protein-containing complex binding, focal adhesion, supramolecular fiber organization (Fig. 3F-H).

PPI interaction analysis was performed for these ECM related DEPs using STRING to obtain a network summarizing their functional associations. A total of 22 proteins formed protein networks (Fig. 4A). Integrins (itga1 and itgb1) formed the hub and had the highest number of connections with collagen (col6a3) and laminin proteins (lama5, lamb1, lama4). Integrin (itga1) also showed interaction with myosin and tropomyosin proteins mapped to actin cytoskeleton. Collagen (col6a3) showed interaction evidence with decorin (dcn) and lumican (lum) (Fig. 4A).

**Figure 4.**
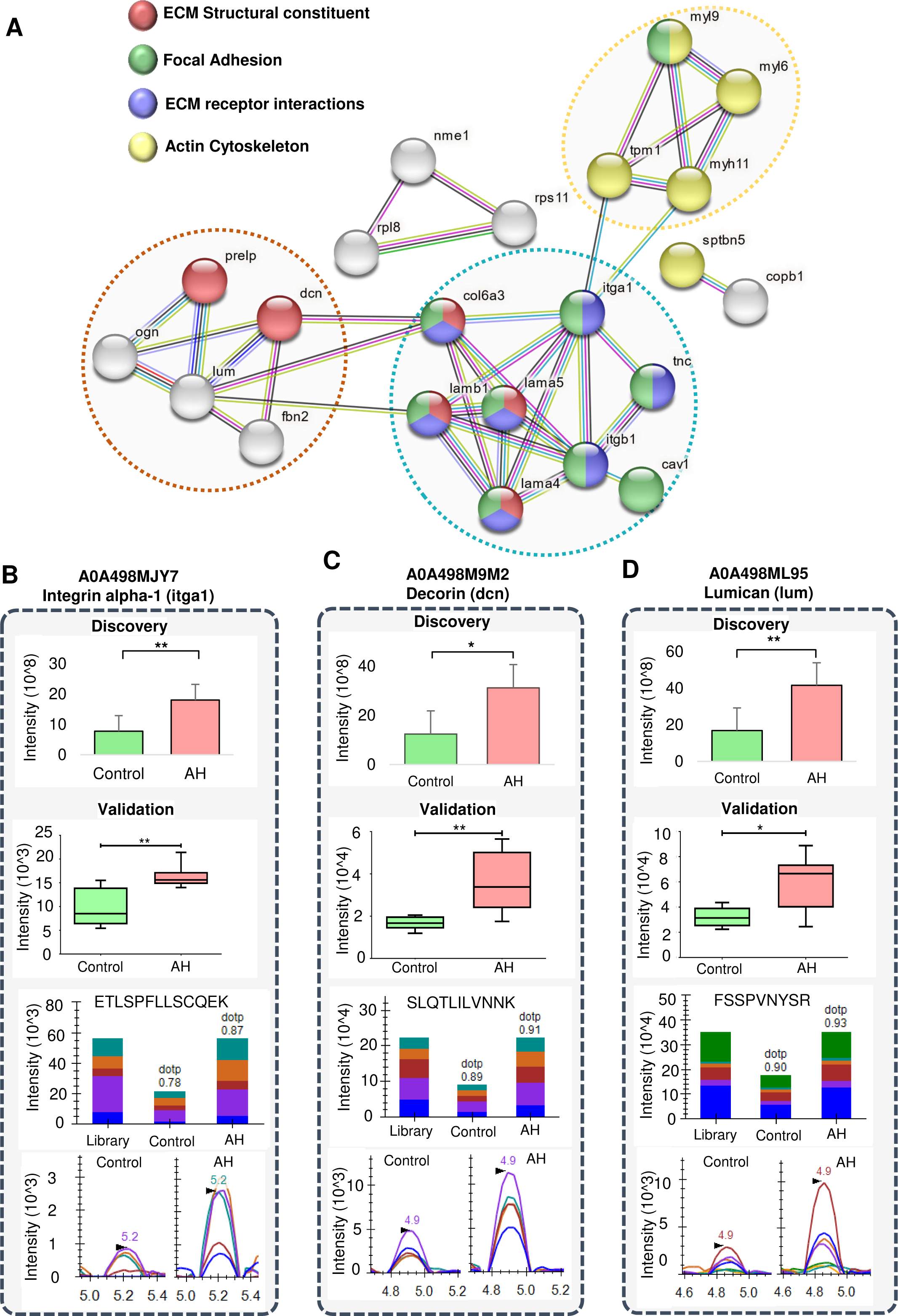
Protein-protein interaction network for ECM related pathways and validation of few candidates using targeted proteomics. **(A)** PPI network for proteins belonging to ECM structural constituent, focal adhesion, ECM receptor interactions, and actin cytoskeleton that showed interactions and three major networks were observed (encircled with dotted line). (**B-D)** Showing differential protein abundance in discovery and targeted experiment for three ECM associated proteins itga1, decorin and lumican, respectively, upregulated in the AH condition (*Ah* infected group) compared to the control condition. (**B-D)** (up to down), the panel represents the protein wise intensities based on the shotgun analysis (DDA abundance) followed by SRM analysis. *p ≤ 0.05 and **p ≤ 0.01. The latter half contains bar plots for peak area and peak view for intensity comparison, for a representative peptide of each protein with AH vs control condition (with dot product (dotp) value based on match with spectral library). Different colors in the bar plots and spectral peaks represent different product ions of the same peptide.

### Targeted proteomic validation using Selected Reaction Monitoring approach

The refinement of SRM validation data in Skyline ended up with 58 peptides corresponding to 10 DEPs. On group comparison of the 6 replicates each from AH and control samples, the overall trend for protein expression was consistent with the DDA data in terms of abundance profile in *Ah* infected (AH condition) samples. Further, applying the criterion of cut off as P value of 0.05 and fold change criterion of 1.5, 25 peptides corresponding to 8 DEPs showed significantly differential peak areas and intensities. Five of these DEPs belong to ECM associated pathways. Three proteins showed significant results with ≥ 2 unique peptides. It included Itga1 (8 peptides), Decorin (6 peptides) and Lumican (2 peptides) showing significantly higher intensity in AH group compared to the Control (Fig. 4B-D). Also, two proteins from focal adhesion pathway namely Mcam (MUC-18 like isoform) and Mfap2 (Microfibrillar associated 2 protein) had one significant upregulated peptide each in the validation data.

The validation experiment included three more proteins which were upregulated in the discovery data. It included two annexin proteins (A0A498LVS6 and A0A498MM94) and GGT5 (Gamma glutamyltransferase–5 like protein) (Table S3). Annexins showed upregulation in AH group in the SRM data each with 3 unique peptides and Ggt5 had one unique significant peptide, and this trend is as par with the discovery data.

## DISCUSSION

Extracellular matrix is well known for its role in immune signalling along with providing structural support to the cell. The diverse effects of the ECM can alter the invasion and dissemination of microbial pathogens in host tissues and greatly influence the overall immune response to infection (20). During tissue damage or infections, it responds by turning on complex biochemical and physical systems that control intercellular communication, which is essential for preserving tissue homeostasis. A scaffold of biochemical and biomechanical signaling is created by the action of several actin cytoskeletal regulators, which are physically coupled to extracellular matrix through integrins (21).

In the current study using proteomic approach, we analysed the overall changes in the gut proteome as a result of *Ah* infection. Several pathways and biological processes including respiratory electron transport, metabolism of vitamins and amide and cell junction organisation were affected. We found the dysregulation of ECM protein components in the gut tissue of host (*Labeo rohita*) during *Ah* infection. Mechanical properties of microenvironments, such as ECM stiffness can modulate the interaction of host cells and pathogen. ECM stiffness-regulated mechanical properties can promote apoptosis and alter the abundance of actin filaments (22) or increases the susceptibility of host cells to infection by bacterial pathogens (23). Highly organised matrix fibres and collagen/elastin cross-linking are among the critical determinants of ECM stiffness (24). In this study, we found increased abundance of elastin (2.7-folds) and Collagen alpha-3(VI) chain (1.7-fold) in AH group. Elevation in the level of Collagen alpha-3(VI) chain has been reported in serum of patients with gastrointestinal disorders (25). It was also found upregulated in intestine tissue of tongue sole *Cynoglossus semilaevis* after *Vibrio vulnificus* infection (26). This might be an indication that *Ah* infection mediates changes in ECM composition that causes matrix rigidity.

To further explore the effect of protein dynamics during *Ah* infection, we looked into the statistically enriched terms/pathways (GO/ KEGG terms) mapped to the significant DEPs, specifically those associated with ECM, actin bundles and focal adhesions. Interestingly, the enriched pathways included Focal adhesions (dre04510) and Integrin cell surface interactions (R-DRE-216083) while actin cytoskeleton (GO:0015629) and extracellular matrix structural constituent (GO:0005201) were among significant GO terms. Focal adhesions (FAs) are macromolecular sites which form mechanical connections between intracellular actin bundles and ECM through clustered integrin receptors. FAs are associated with immune system and play an important role during infections or diseases. It has been reported that pathogens exploit focal adhesions to ensure their uptake, survival, and dissemination (27). They serve as scaffolds for many signalling pathways triggered by cell surface adhesion molecules, receptors as integrin or a mechanical tension on the cell.

Integrins are alphabeta heterodimeric integral proteins and members of the cell adhesion receptor superfamily expressed in all metazoans. They are referred as the key molecules in cell-microenvironment and cell-cell communication. It was reported that expression of integrin beta was increased in *Penaeus monodon* after challenging with bacteria *Vibrio harveyi* and *V. anguillarum.* Further the authors reported that inhibition of integrin beta gene caused downregulation of innate immunity related genes (28). We also found increased abundance of integrin beta 1 (3.3-fold) and integrin alpha 1 (2.3-fold) in the AH group, in the discovery and the validation experiment. Another important protein from the Integrin cell surface interactions pathway was Tetraspanin (tspan) which showed 2.9 folds increase during infection in our study. Tetraspanins represent a conserved superfamily of four-span membrane proteins, which are associated with innate immune response and bacterial adhesion as well (29). In a study on giant freshwater prawn, *Macrobrachium rosenbergii* infected with *Ah*, the expression of tspan8 (Tetraspanin 8) was increased in hepatopancreas and gill tissue. According to the authors, pre-incubation of peptides from the long extracellular loop of tspan8 protein decreased the apoptosis caused by *Ah* infection in prawn tissue (30). Such results indicate that integrins and tspan have important role in host response to *Ah* infection.

Subunits of myosin (myl6, myl9, myo1c), tropomyosin (tpm1), phosphoglucomutase (pgm5), a protein from tubulin polymerisation promoting protein family (tppp3), lumican (lum) and a few more proteins showed increased abundance in AH group and mapped to actin cytoskeleton (GO:0015629). These proteins control the cytoskeletal reorganisation during infection and leads to changes in cell behaviour. Latent-transforming growth factor beta-binding 1-like isoform X1 (ltbp1) which is 1.8-fold increased during *Ah* infection was also mapped to actin cytoskeleton. It has been reported that LTBP-1 forms a complex facilitated by disulfide linkage with TGFβ propeptide (known as LAP or latency associated peptide) in the endoplasmic reticulum before secretion. This binding assists in proper folding and release of large latent complex and its interaction with ECM. Transforming growth factor (TGF-β) is a crucial immunoregulatory cytokine that regulates cell proliferation, differentiation, survival, migration, and apoptosis in both healthy and pathological circumstances. TGF-β1 has been found upregulated with poly I:C or LPS induction in *Culter alburnus*, a teleost (31). LTBP1 is reported to interact with fibrillin (and/or fibronectin) and other matrix components which helps in release of TGF-β from the latent complex to bind to its receptor (32). We identified Fibrillin-2-like isoform X1 (fbn2) in this study, a structural constituent of extracellular matrix, mapped to focal adhesion and showed a 2.9-fold increased abundance during infection.

It has been reported that the first and the most important step towards the successful establishment of infection is the adherence of bacteria to the host cells. One crucial tactic is their interaction with host ECM proteins like fibronectin, collagen, elastin, vitronectin and laminin (33). We found increased abundance of laminin proteins viz; laminin subunit alpha-4 (1.8-fold), laminin subunit alpha-5 (1.8-folds) and laminin subunit beta-1 (1.7-folds), elastin (2.7-folds) and collagen alpha-3(VI) (1.7-folds). Laminins interact with integrins, collagens and other ECM components that allow the assembly and integrity of basement membrane. Laminin interaction with other ECMs also influence the migration and the function of immune cells (34). Another protein, galectin 3 (lgals3) mapped to focal adhesion pathway showed increased abundance in AH group. Galectin 3 is a glycan binding lectin, reported to function in host defence as an opsonin. Galectin 3 when bound to LPS, promoted the bacterial phagocytosis by microglia i.e., brain macrophages (35). The dysregulation of these ECM proteins in the study indicates these candidates might be important players for pathogen invasion during *Ah* infection (Fig. 5A).

**Figure 5.**
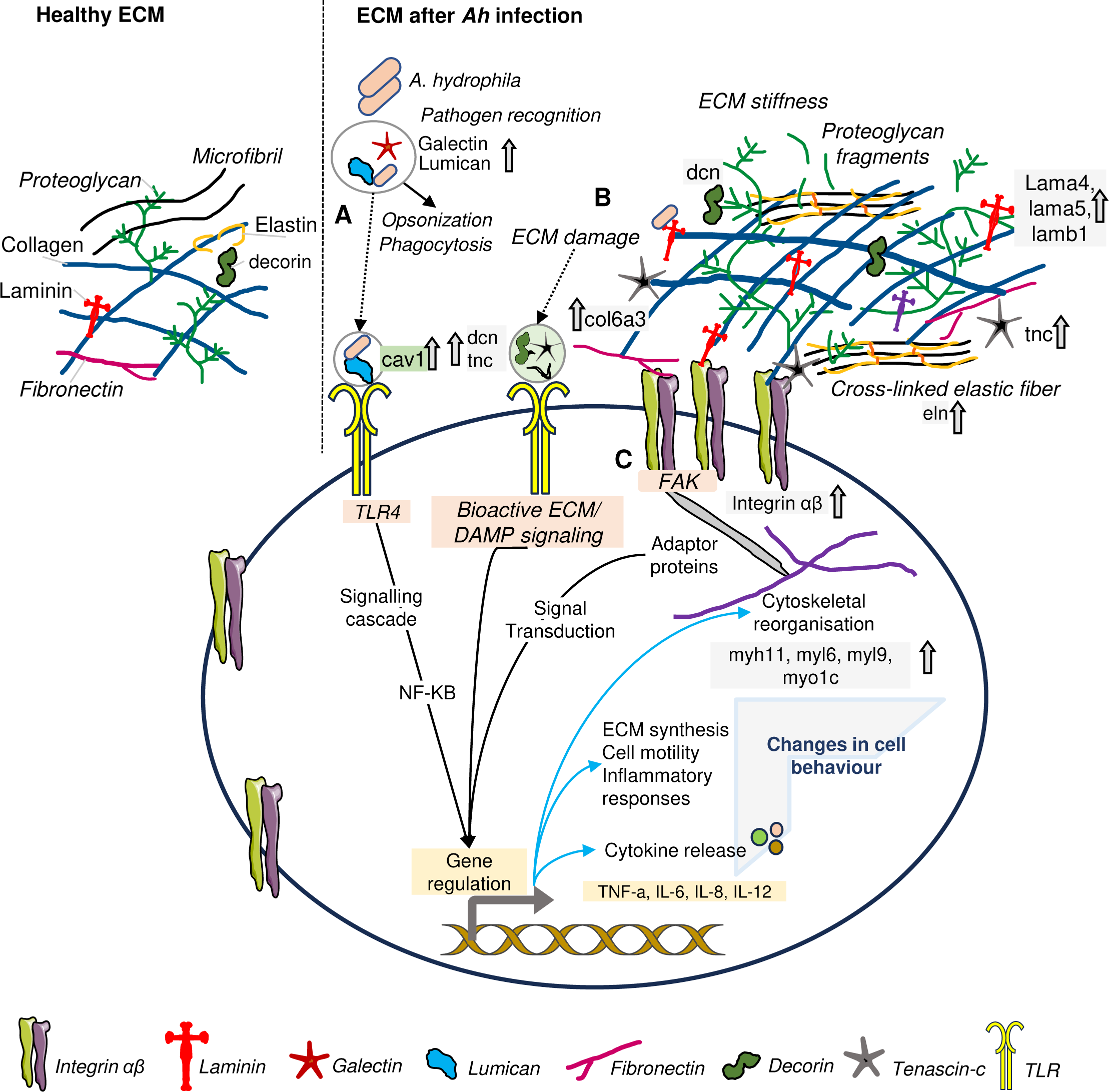
Hypothesised interplay of *bacteria* and host tissue during *Ah* infection: Left side shows the healthy ECM with normal architecture of ECM components and, right side shows the compact and stiff ECM during *Ah* infection with cross-linked fibres and ECM fragments. **(A)** Interaction of the bacteria with the ECM proteins like fribronectin, galectin, lumican, laminin, helps in pathogen recognition and adherence with the host cells. These ECM proteins also act as opsonins to promote opsonisation and phagocytosis. These proteins are also linked with TLR receptor mediated cell response to affect gene regulation. **(B)** Once the pathogen is entered, it degrades the ECM through matrix degrading enzymes resulting into bioactive ECM fragments. Bioactive ECM fragments and DAMP immunomodulators as decorin (dcn), tenascin-c (tnc), lumican further start cascade of signals by interacting with fibroblasts or surface receptors like TLRs on immune cells. As a result, more ECM proteins are produced and cross-linked ECM fibres are formed leading to stiffness. **(C)** Increased rigidity and stiffness in ECM followed by larger and stable focal adhesions lead to signaling through focal adhesion kinase and integrins. Such processes will further initiate mechanotransduction ending into inflammatory responses, release of cytokines, cytoskeletal reorganisation and changes in cell behaviour for pathogen clearance and tissue repair. [Upside arrow represents upregulated proteins in this study.

In an earlier study, the transcriptional level of tenascin-c was found increased in ulcerated areas during inflammatory bowel disease in murine model of IBD (36). Tenascin-C is a pro-inflammatory ECM protein, is upregulated with tissue injury and cellular stress. Tenascin-C interacts with receptors like TLR4, integrins to initiate immunomodulatory effects. IThe subsequent interaction with immune cells, results in the induction of soluble proinflammatory mediators, such as interleukin (IL-6, IL-8, IL-1β and IL-18), and tumor necrosis factor (TNF) (36). Based on these findings and our results, it could be anticipated that increased abundance of tenascin-C during *Ah* infection would lead to an increase in proinflammatory cytokines and chemokines, coordinating the activation and recruitment of innate immune cells (Fig. 5B).

Once the pathogen has invaded and caused degradation in ECM, ECM fragments may be released to serve as “danger signals” i.e., damage associated molecular patterns (DAMPs) during infection, inflammation, injury, or even aging. Such immunoregulatory fragments can be produced when the ECM is broken down by matrix-degrading enzymes (37). They lead a cascade of signalling which results in initiation of pathogen clearance and tissue repair. In this study, AH group was found with higher abundance of few proteins, reported to act directly or indirectly as immunomodulators. Such proteins include decorin (2.5-fold), lumican (2.5-fold), Tenascin-like isoform X1 (2.1-fold) and fibrillin-2 (2.9-fold). These ECM proteins are linked with actin cytoskeleton and focal adhesion. Decorin is a small leucine-rich repeat proteoglycan, which coordinates pro- and anti-inflammatory cytokines and macrophage recruitment by directly binding to TLR2 and TLR4 receptors. Lumican is similar to decorin and known to bind with LPS and coordinates the innate immune response by enhancing LPS/TLR4 mediated proinflammatory responses (38). It has been reported that the circulating level of lumican was increased in human and mouse during sepsis and mice lacking lumican showed poor bacterial clearance. The author also reported that a protein caveolin (cav1) is required to maintain the lumican on cell surfaces (39). In this study, Caveolin was increased by 1.6-fold in AH group and mapped to the focal adhesion pathway.

With these immunomodulators, the signalling event starts by interacting with pattern recognition receptors (PRRs) as Toll-like receptor present on immune cells like dendritic cells and macrophages, other cells as epithelial cells and fibroblasts (40) (27). As a result, more ECM proteins are produced and cross-linked ECM fibres are formed (Fig. 5B-C). It results in increased rigidity and stiffness in ECM followed by larger and stable focal adhesions leading to focal adhesion signalling and changes in cell behaviour. Overall, the results indicate these ECM proteins have an important role during *Ah* infection as they coordinate initiation of immune response during *Ah* infection through damage signals and their interaction with surface receptors as TLRs and focal adhesion signaling.

## CONCLUSIONS

The investigation into the proteomic profile of the gut tissue in *Labeo rohita* following infection with *Aeromonas hydrophila (Ah)* reveals substantial alterations in the extracellular matrix (ECM) proteome. Notably, 39 proteins associated with Integrin cell surface interactions, focal adhesion, actin cytoskeleton, and extracellular matrix structural organization exhibit differential expression during *Ah* infection. This observation underscores the intricate interplay between the host’s immune system and the ECM. The findings suggest a reciprocal relationship, wherein signals from the ECM play a crucial role in coordinating immune responses. Additionally, immune cells contribute to ECM repair and regeneration through the release of cytokines, influencing ECM synthesis, cytoskeleton reorganization, and alterations in cell behavior. To our best knowledge, this is the first study to present landscape of gut proteome during *Ah* infection in this food fish. Our findings open avenues for future research in the field of aquaculture and disease control in *Labeo rohita* and related carps. Further investigations could focus on the specific mechanisms through which dysregulation of ECM proteins influences immune responses and host-pathogen interactions. Understanding the molecular signaling pathways during *Ah* infection could provide targets for therapeutic interventions.

## ACKNOWLEDGEMENTS

This work was supported by the Department of Biotechnology (BT/PR15285/AAQ/3/753/2015) Govt. of India to S.S and M.G. M.N was supported by University Grants Commission (UGC). We acknowledge ICAR-Central Institute of Fisheries Education, Mumbai for supporting this work. We acknowledge MASS-FIITB at IIT Bombay supported by the Department of Biotechnology (BT/PR13114/INF/22/206/2015) for Mass-spectrometric data acquisition.

## DATA AVAILABILITY

The protein database (.FASTA) and raw mass spectrometry data (.raw) have been deposited to the ProteomeXchange Consortium via the PRIDE partner repository. All result output files for protein identification are also submitted in text (.text) format along with the parameter file. Also, the spectral library (.blib) generated using the discovery data for analysing the targeted data is uploaded. The identifier PXD037850 can be used to retrieve all of the data. (Reviewer account details: Username: Password: UaWQZTvo) The transition lists, skyline documents, and all SRM raw (.raw) data for the Selected reaction monitoring (SRM) experiment have been submitted to Panorama public that can be accessed through the given link https://panoramaweb.org/rohugutproteomicsah.url (Reviewer account details: Password: gxyJdlEl).

## AUTHOR’S CONTRIBUTIONS

**Mehar Un Nissa:** Conceptualization, Methodology, Investigation, Validation, Formal analysis, Writing - Original Draft, Review and Editing, Visualization: All original data

**Nevil pinto**: Resources, Formal analysis, Writing - Original Draft, Visualization

**Biplab Ghosh**: Formal analysis, Visualization, Writing - Original Draft, Review and Editing

**Anwesha Banerjee**: Formal analysis, Visualization, Writing - Original Draft, Review and Editing

**Urvi Singh**: Formal analysis, Visualization, Writing - Original Draft, Review and Editing

**Mukunda Goswami**: Resources, Conceptualization, Writing – Original Draft, Review & Editing

**Sanjeeva Srivastava**: Supervision, Resources, Conceptualization, Writing - Original Draft, Review & Editing

## SUPPLEMENTAL INFORMATION

This article contains three supplementary Tables (Table S1 to S3) (.xls).

## Supplementary Tables

**Table S1**| Details of discovery data (DDA) used for LFQ analysis

**Table S2**| Details of pathways, gene ontology and protein-protein interaction data analysis using Metascape and STRING

**Table S3**| Details of SRM data used for targeted proteomic analysis

